# Harringtonine has the effects of double blocking SARS-CoV-2 membrane fusion

**DOI:** 10.1101/2022.01.22.477323

**Authors:** Shiling Hu, Nan Wang, Shaohong Chen, Qiang Ding, Cheng Wang, Weina Ma, Xinghai Zhang, Yan Wu, Yanni Lv, Zhuoyin Xue, Haoyun Bai, Shuai Ge, Huaizhen He, Wen Lu, Tao Zhang, Yuanyuan Ding, Rui Liu, Shengli Han, Yingzhuan Zhan, Guanqun Zhan, Zengjun Guo, Yongjing Zhang, Jiayu Lu, Jiapan Gao, Qianqian Jia, Yuejin Wang, Hongliang Wang, Shemin Lu, Huajun Zhang, Langchong He

**Author notes:** Correspondence Author: Langchong He and Huajun Zhang, **Address for correspondence**, School of Pharmacy, Xi’an Jiaotong University, Yanta Westroad, Xi’an 710061, China., State Key Laboratory of Virology, Wuhan Institute of Virology, Center for Biosafety Mega-Science, Chinese Academy of Sciences, Wuhan 430071, China. These authors contributed equally to this work (Shiling Hu, Nan Wang, Shaohong Chen, Qiang Ding, Huajun Zhang, Langchong He).

## Abstract

Fusion with host cell membrane is the main mechanism of infection of SARS-CoV-2. Here, we propose a new strategy to double block SARS-CoV-2 membrane fusion by using Harringtonine (HT), a small-molecule antagonist. By using cell membrane chromatography (CMC), we found that HT specifically targeted the SARS-CoV-2 S protein and host cell TMPRSS2, and then confirmed that HT can inhibit pseudotyped virus membrane fusion. Furthermore, HT successfully blocked SARS-CoV-2 infection, especially in the delta and Omicron mutant. Since HT is a small-molecule antagonist, it is minimally affected by the continuous variation of SARS-CoV-2. Our findings show that HT is a potential small-molecule antagonist with a new mechanism of action against SARS-CoV-2 infection, and thus HT mainly targets the S protein, and thus, greatly reduces the damage of the S protein’s autotoxicity to the organ system, has promising advantages in the clinical treatment of COVID-19.

## Results and discussion

Since the outbreak of SARS-CoV-2 infection in December 2019, it has still wreaked havoc worldwide (**Fig. 1a, b**). Currently, many types of vaccines have been widely used for the prevention of SARS-CoV-2 infection^1^, while only two anti-COVID-19 drugs have been approved in the world^2,3^. These drugs target the replication process after SARS-CoV-2 enters host cells^4,5^. Given the continuous variation of SARS-CoV-2 into mutants such as Delta and Omicron mutants, effective therapeutic drugs that are capable to treat mutant infections are an urgent demand in the clinic.

**Fig. 1.**
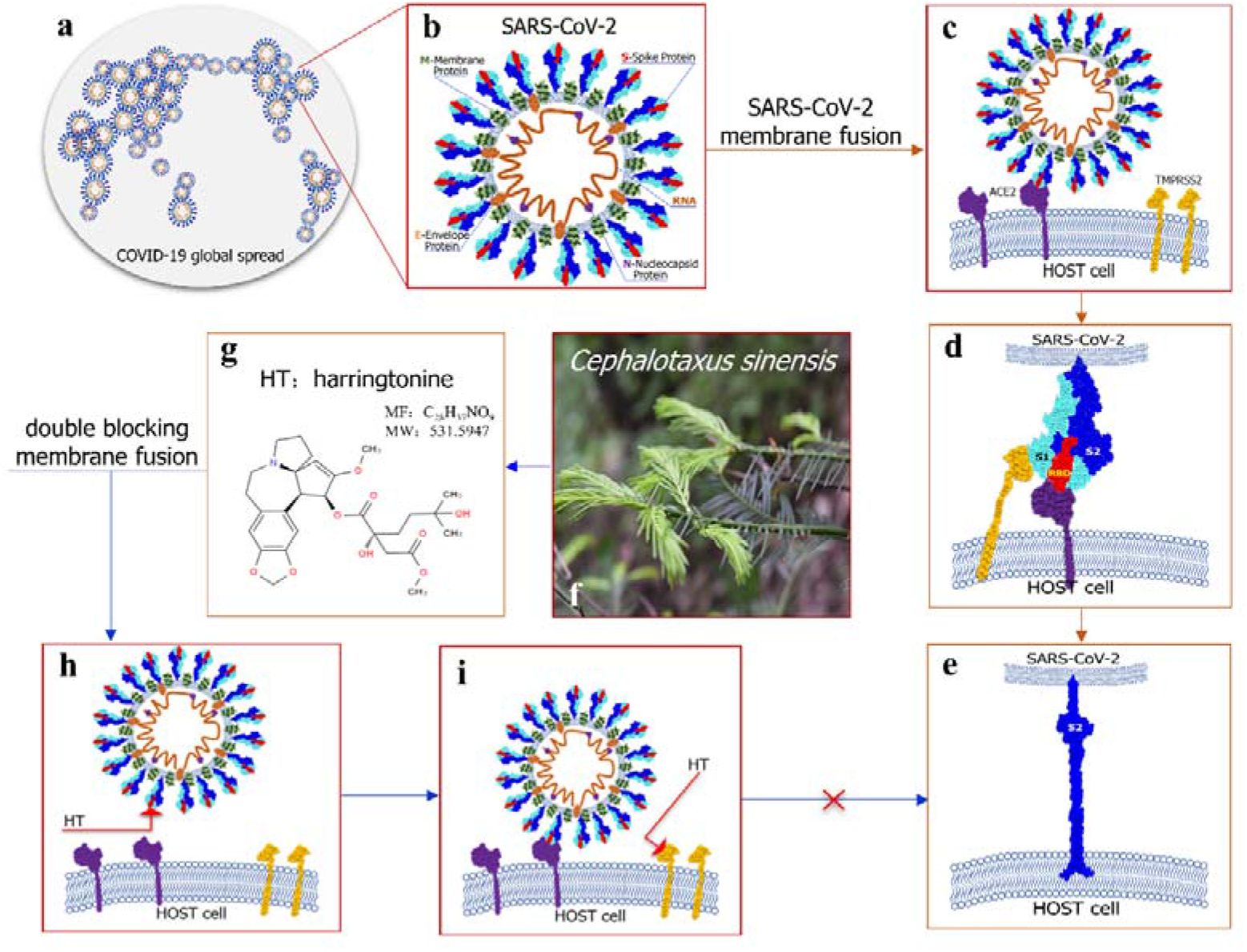
A new strategy of double blocking SARS-CoV-2 membrane fusion using harringtonine (HT). **a**, The COVID-19 pandemic is spreading globally. **b**, SARS-CoV-2 is mainly composed of the spike (S), membrane (M), and envelope (E) proteins. There is a viral RNA chain of the nucleocapsid (N) protein inside SARS-CoV-2. **c-e**, SARS-CoV-2 membrane fusion process includes the formation of the S-ACE2 complex, which is hydrolysed by TMPRSS2 and S2 fusion peptide anchored to the host membrane. **f-g**, HT was isolated from *Cephalotaxus sinensis*. **h-i**, Our aim is to find an antagonist such as HT that blocks SARS-CoV-2 membrane fusion by simultaneously targeting S protein and the host TMPRSS2.

Natural medicinal plants are an important source of small molecule new drugs^6^. The *Cephalotaxus sinensis* is a unique medicinal plant from Qinling Mountains of China^7^, and the average content of HT is 0.0045% (**Fig 1. f-g**). In this study, we propose a new strategy of treatment to be directed against SARS-CoV-2 membrane fusion through the S protein (**Fig.1c-e**). HT, a small molecule antagonist, targets the S protein to inhibit its binding to the ACE2 receptor in host cells; simultaneously, it targets the host cell TMPRSS2 against the enzymatic hydrolysis of S protein (**Fig1. h-i**). Our experiments showed that HT is an effective antagonist and acts by double blocking SARS-CoV-2 membrane fusion.

### 1. HT Targeted the S protein of SARS-CoV-2, and to TMPRSS2 in host cells

Cell membrane chromatography (CMC) is a biomimetic analysis technology used to study interactions between drugs and their receptors^8^. In this study, the S/CMC and TMPRSS2/CMC models were established to test the affinities between HT and S protein as well as TMPRSS2^9^. The reversibility of HT on the S protein and TMPRSS2 was determined. In addition, the binding modes of HT were simulated by computer molecular docking technology.

The *K*_D_ values of HT were 1.36 ± 0.10 ΔM (S protein) and 15 ± 3.34 ΔM (TMPRSS2) ^10^. The A_S_ values were 1.71 (S protein) and 5.27(TMPRSS2) (**Fig. 2a-d**). CMD showed that HT can form hydrogen bonds with the GLY696, SER494, TYR449, and ARG403 residues in the RBD domain of the S protein (PDB ID: 6M0J), and these sites were key binding sites between the S protein and ACE2. HT can also bind the SER441, GLY462, and GLY464 residues in the domain of TMPRSS2 (PDB ID: 7MEQ) (**Fig. 2e-f**). These findings show that HT has a strong affinity for SARS-CoV-2 S protein and TMPRSS2, and also shows some irreversibility, which is very beneficial for blocking SARS-CoV-2 membrane fusion.

**Fig. 2.**
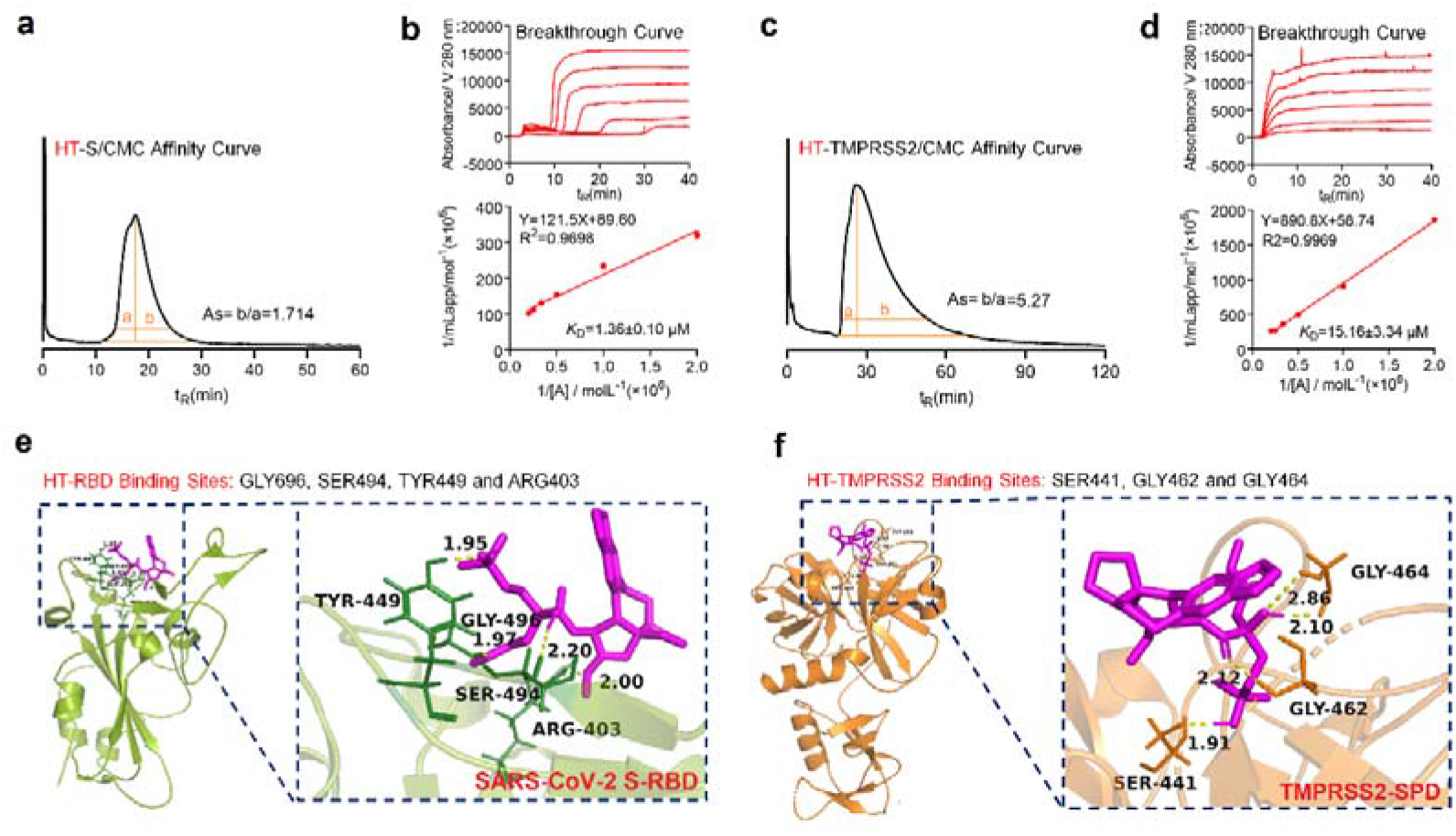
HT specifically targets the S protein of SARS-CoV-2 and host cell TMPRSS2. **a-b**, Affinity and breakthrough curves of HT were obtained using the S/CMC-HPLC method and **c-d**, the TMPRSS2/CMC-HPLC method. **e-f**, Computer molecular docking (CMD) was used to identify the hydrogen bonds between HT and the receptor-binding domain (RBD) of S protein, and those between HT and the serine protease domain (SPD) of TMPRSS2. Graphs display mean±S.E.M. One-way ANOVA (and nonparametric or mixed) (\**P*<0.05, \*\**P*<0.01, \*\*\**P*<0.001 *vs*. 0)

### 2. HT inhibits pseudotyped virus membrane fusion

The interaction between ACE2 and the S protein plays a key role in SARS-CoV-2 infection^11^. We used NanoBiT® to evaluate the effect of HT on the interaction between ACE2 and The S protein **(Fig. 3a)**. As shown in Fig 3b, HT can dose-dependently inhibit cell-to-cell membrane fusion induced by the binding of ACE2 and S proteins (S495) **(Fig. 3b-c, and e)**. In addition, HT also effectively inhibited ACE2^h^ cell membrane fusion induced by Delta virus mutated S protein-L452R, T478K, and P681R sites (S605) **(Fig. 3b, d, and f)**. It was evident from the micrographs that HT dose-dependently inhibited S495 and S605-induced syncytial formation.

**Fig. 3.**
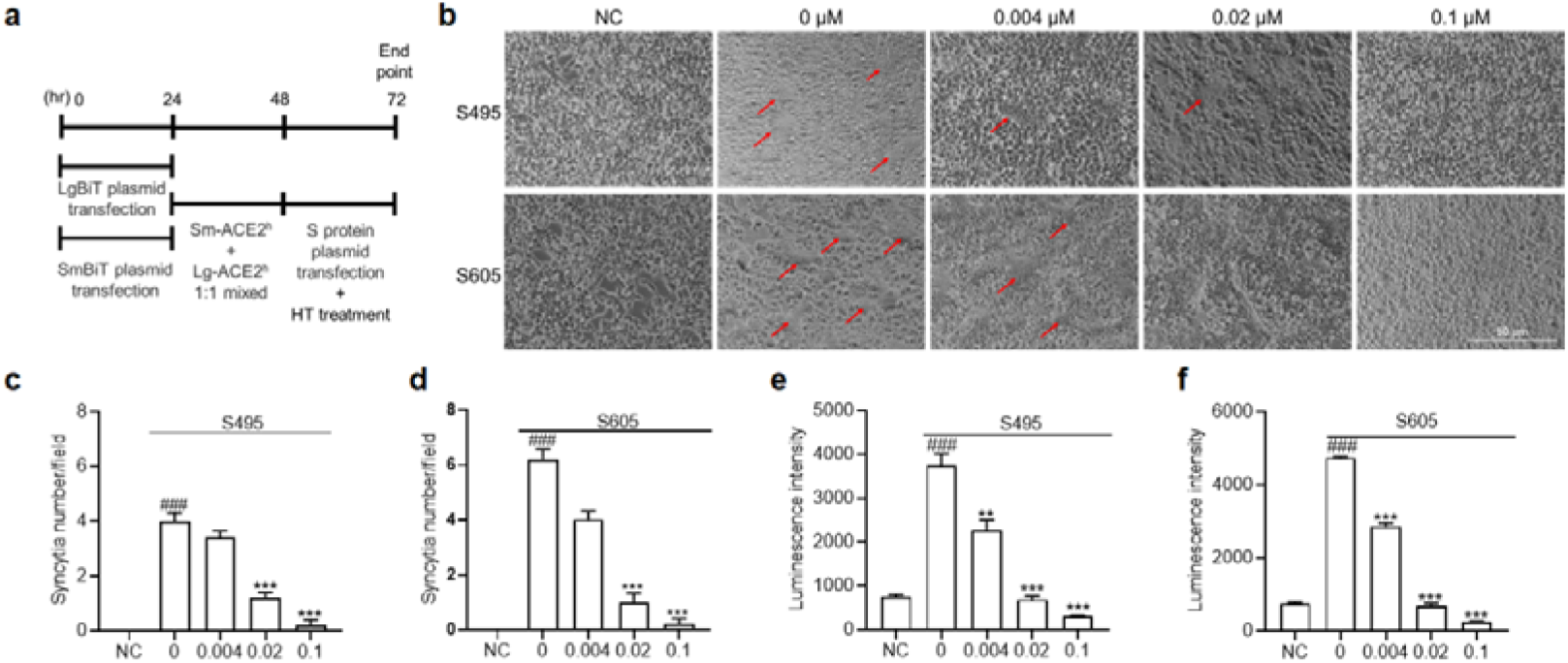
HT simultaneously membrane fusion through S protein. **a**, Protocol of ACE2h cells membrane fusion assay; ACE2h cells were transfected with LgBit plasmid and SmBit plasmid respectively for 24h.Then, 1:1 mixed and incubated for the night. After that, mixed cells were transfected with S protein plasmid for 24h and the cells were collected and used to evaluate Luminescence intensity. **b**, HT inhibited S495 and S605 induced ACE2h cells membrane fusion. **c-d**, Quantitation of numbers of syncytia in cells as described in (b). Values are syncytia number per microscope field. **e-f**, Luciferase were measured to indicate the effect of HT to S495 and S605-induced syncytial formation. Graphs display mean ± S.E.M. One-way ANOVA (and nonparametric or mixed) (\**P*<0.05, \*\**P*<0.01, \*\*\**P*<0.001 *vs*. 0)

We used a pseudotyped virus, it can be used to evaluate the antiviral effects of drugs only in a P2 Lab^12,13^**(Fig. 4a)**. The pseudovirus has a significant infection advantage for ACE2^h^ cells^14^. ACE2^h^/TMPRSS2^h^ cells were successfully established by stably overexpressing TMPRSS2 in ACE2^h^ cells^15^. The pseudo-type entry rates in ACE2^h^/TMPRSS2^h^ cells were greatly improved compared with those in ACE2^h^ cells when pseudotyped viruses were used to infect different cells. The ACE2^h^/TMPRSS2^h^ cell model works well because the Camostat (TMPRSS2 antagonist) considered to be an effective antagonist of TMPRSS2 and can effectively inhibit pseudovirus infection (**Fig. 4b**).

**Fig.4.**
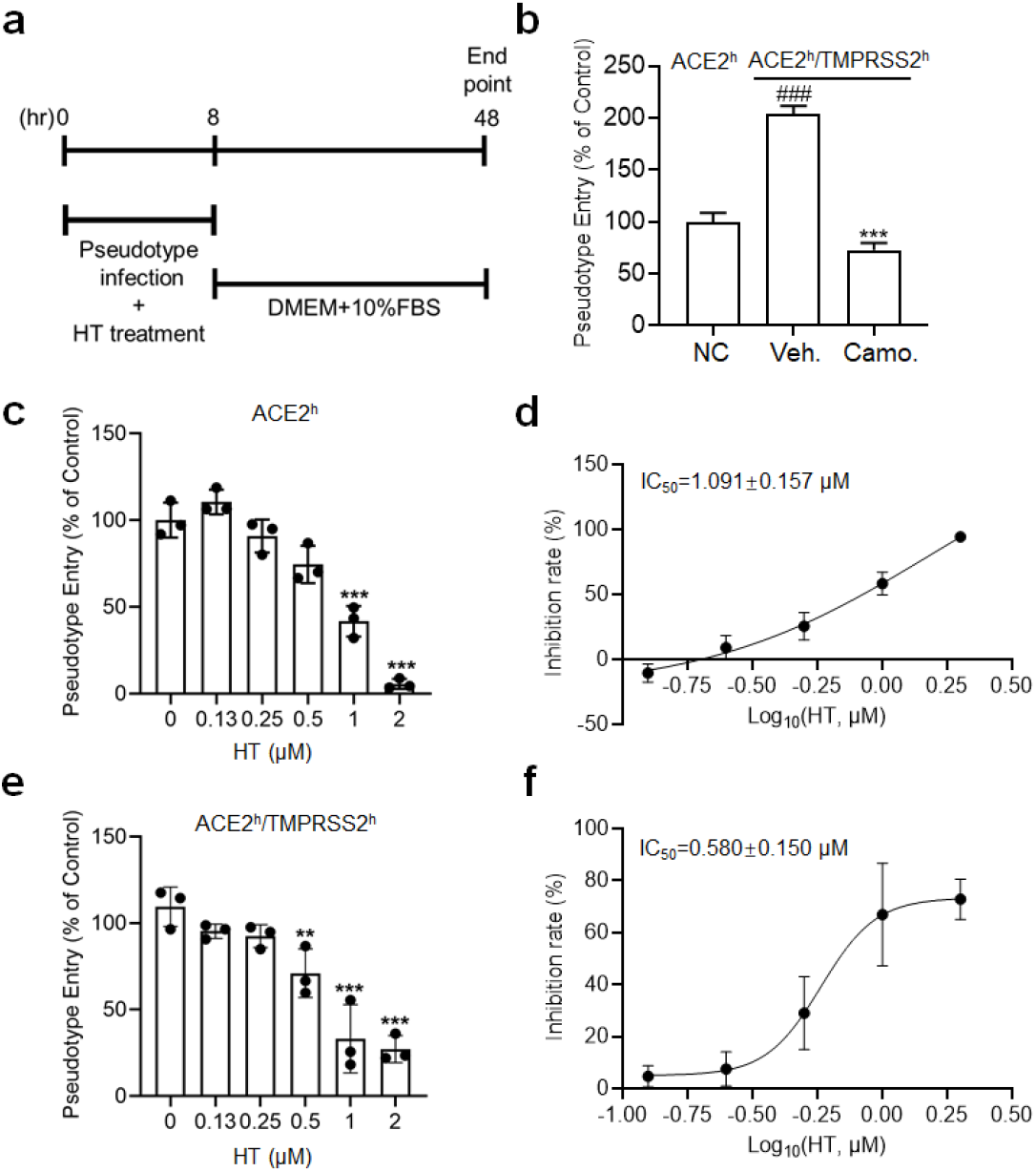
Effect of HT on the entrance of SARS-CoV-2 spike pseudotyped virus into ACE2^h^ and ACE2^h^/TMPRSS2^h^ cells. **a**, Protocol of pseudotyped virus entry assay; ACE2^h^ cells were pretreated with pseudotyped SARS-CoV-2 virus combined with different doses of HT for 8 h. Then, the cells were transferred to fresh cell culture medium and incubated for 48 h; the cells were collected and used to evaluate pseudotype entry rates. **b**, TMPRSS2 was transfected into ACE2^h^ cells (ACE2^h^/TMPRSS2^h^ cell), which were then infected with pseudotype combined with vehicle (Veh.) or camostat (Cam.) The entrance of pseudotype virus into ACE2^h^/TMPRSS2^h^ cells was inhibited by camostat. **c-d**, HT inhibited pseudotyped virus entry into ACE2^h^ cells. **e-f**, HT inhibited pseudotyped virus entry into ACE2^h^/TMPRSS2^h^ cells dose-dependently. Graphs display mean±S.E.M. One-way ANOVA (and nonparametric or mixed) (\**P*<0.05, \*\**P*<0.01, \*\*\**P*<0.001 *vs*. 0)

The TMPRSS2 enzyme on host cells plays a key role in pseudovirus virus membrane fusion. HT dose-dependently inhibited pseudovirus infection in ACE2^h^ and ACE2^h^/TMPRSS2^h^ cells, with IC_50_ values of 1.091 ± 0.157 ΔM and 0.580 ± 0.15 ΔM, respectively. The latter is obviously stronger than the former (**Fig. 4c-f**). The results show that HT is an effective antagonist against SARS-CoV-2 infection, and TMPRSS2 has a key synergistic effect.

### 3. HT as an antagonist for anti-SARS-CoV-2 infection

The anti-SARS-CoV-2 effects of HT in vitro were investigated in SARS-CoV-2ΔN trVLP with N protein deficient^16^, SARS-CoV-2-WIV04 original strain, and SARS-CoV-2-Delta 630 mutant strain.

The survival rate of Caco-2-N cells was more than 70% under 1 ΔM HT treatment (**Fig. 5a**). HT dose-dependently decreased SARS-CoV-2ΔN trVLP infection in Caco-2-N cells, with an IC_50_ value of 0.125 ± 0.052 ΔM (**Fig. 5b-d**).

**Fig. 5.**
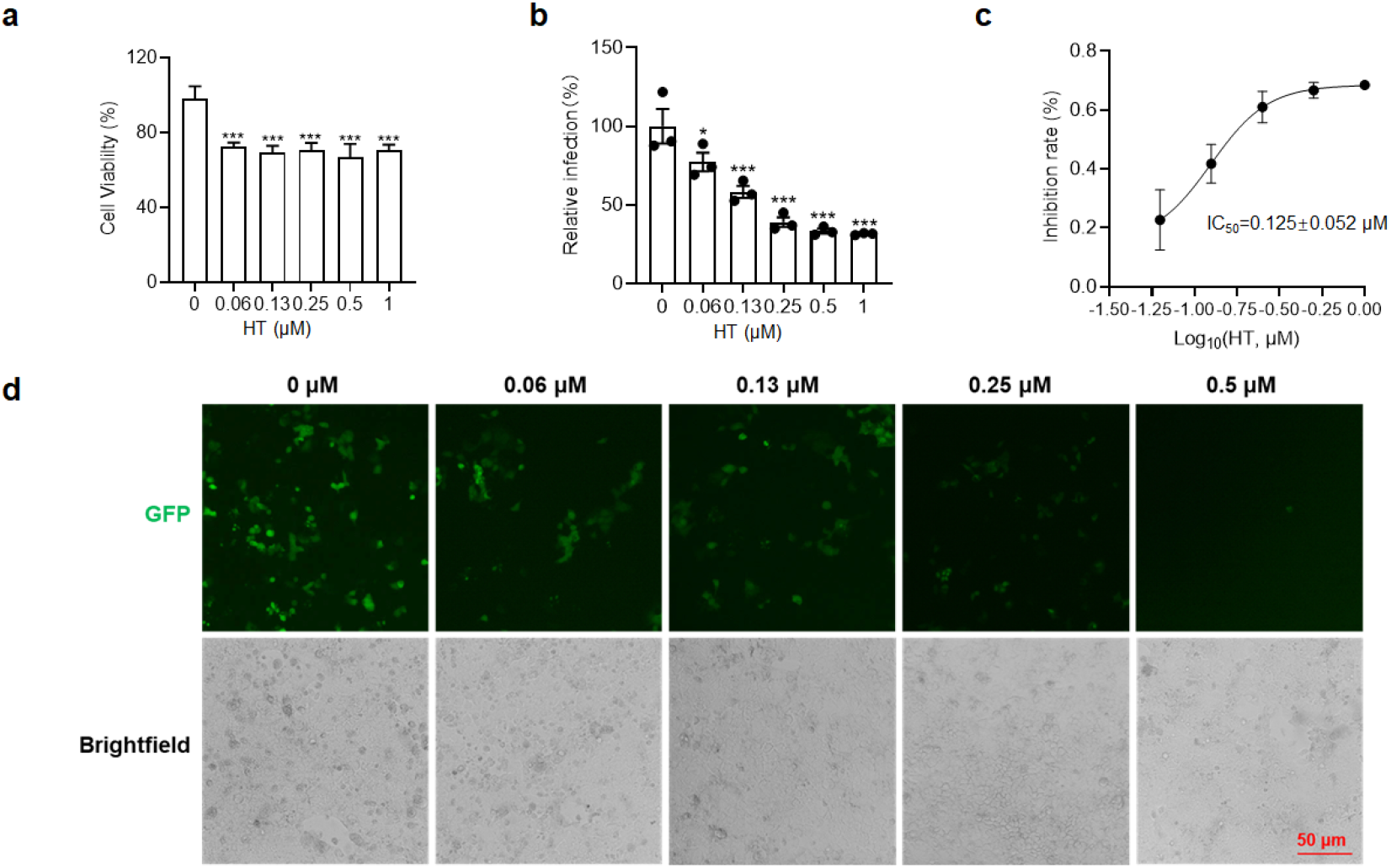
HT successfully blocks SARS-CoV-2/ΔN infecting Caco 2-N cells. **a**, Viability of Caco2-N cells treated with 1 ΔM HT for 72 h. **b-c**, HT in the range of 0.06~1 ΔM dose-dependently decreased SARS-CoV-2ΔN trVLP infection in Caco-2-N cells. **d**, Representative image of HT inhibiting SARS-CoV-2/ΔN infecting Caco 2-N cells. Graphs display mean ± S.E.M. One-way ANOVA (and nonparametric or mixed) (\**P*<0.05, \*\**P*<0.01, \*\*\**P*<0.001 *vs*. 0)

SARS-CoV-2-WIV04 has been one of the original strains since the outbreak of COVID-19. Vero-E6 cells are African green monkey kidney cell lines that highly express the ACE2 receptor^17^. In Vero E6, HT dose-dependently inhibited the cytopathic effect CPE caused by WIV04 virus (**Fig. 6a**). In Vero E6 cells, the CC_50_ and IC_50_ values of HT were 1.35 ΔM and 0.217 ΔM, respectively, and the selectivity index (SI=CC_50_/EC_50_) was 6.22 (**Fig. 6b-c**). Repeated epidemics are caused by the constant mutation of the SARS-CoV-2. The delta mutant strain was found in India as early as October 2020^18,19^ rapidly spreading and became a major concern in the containment of the pandemic^20,21^. On November 9, 2021, the first Omicron sequence available was from a specimen collected in Botswana. Ever since the identification of Omicron, the variant appears to rapidly spread. At the time of this writing, Omicron has become a major concern in the containment of the current pandemic^22,23^. The same evaluation tests for HT were performed on the Delta 630 and Omicron mutant. The results further confirmed that HT inhibit SARS-CoV-2-Delta 630 virus and Omicron infections at a lower inhibitory dose of 0.074 ΔM and 0.025 ΔM (**Fig. 6d-e**). The IC_50_ value of HT was as low as 0.101 ΔM and 0.042 ΔM, with a SI value of 13.4 and 32.1, which was almost five times that of the WIV04 original strain (**Fig. 6f-g**). With the increase of mutation sites on RBD, the antiviral effect of HT was significantly enhanced rather than decreased (**Fig. 6h**).

**Fig. 6.**
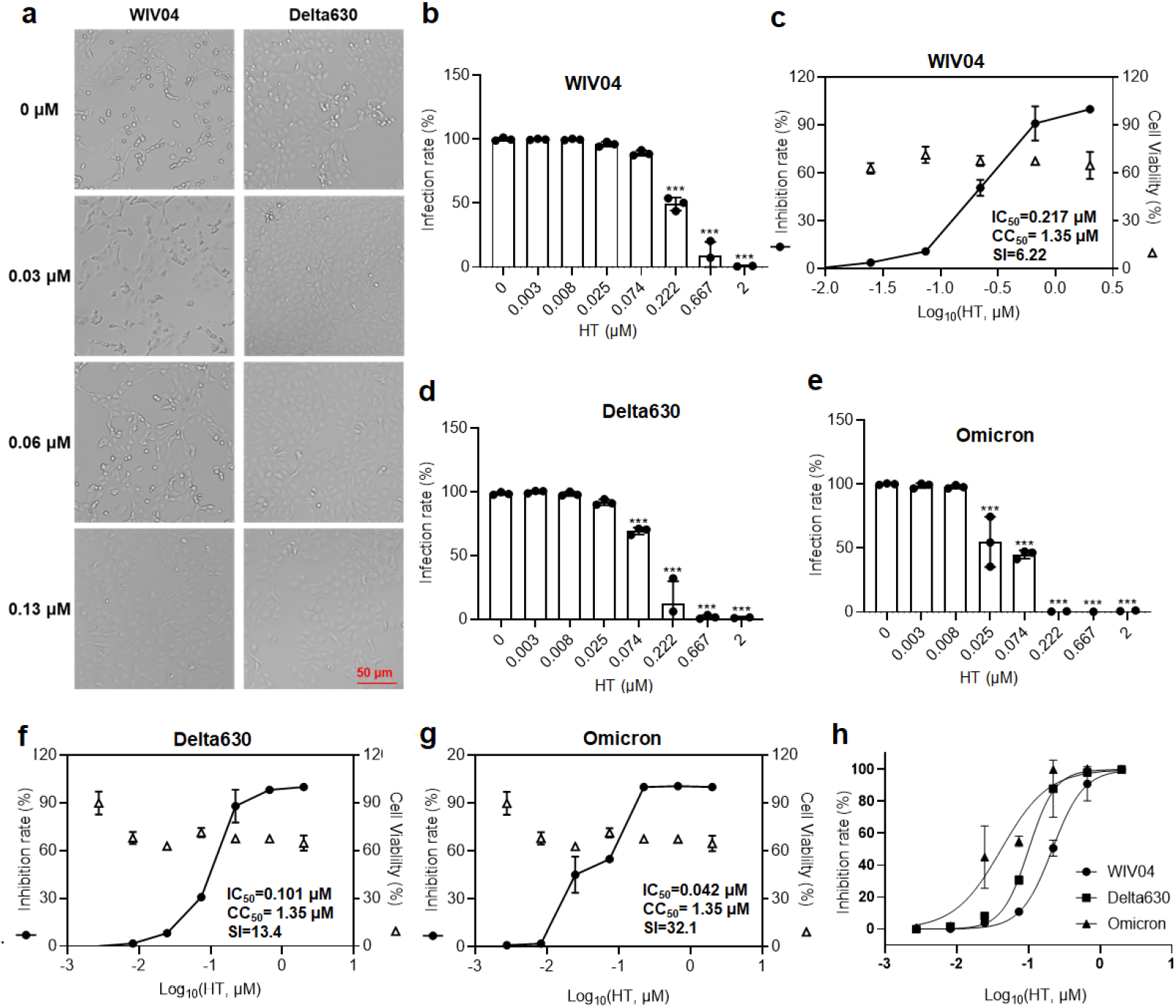
HT successfully blocks SARS-CoV-2 infection. **a**, Representative image of HT inhibiting SARS-CoV-2-WIV04 and SARS-CoV-2-Delta 630 virus-induced cytopathic effects (CPE). **b-c**, Real-time RT-PCR showed that HT in the range of 0.03~2 ΔM dose-dependently inhibited SARS-CoV-2-WIV04 virus infection in Vero E6 cells. **d-e**, Real-time RT-PCR showed that HT significantly inhibited SARS-CoV-2-Delta 630 and Omicron virus infection in Vero-E6 cells, with a lower inhibiting dose. **f-g**, The IC_50_, CC_50_ and SI of HT on the two variant strains of SARS-CoV-2. **h**, HT inhibits SARS-CoV-2 and its variant strains from infecting Vero E6 cells. Graphs display mean ± S.E.M. One-way ANOVA (and nonparametric or mixed) (\**P*<0.05, \*\**P*<0.01, \*\*\**P*<0.001 *vs*. 0)

In line with the viral immunofluorescence, the expression of S protein was dramatically decreased in those cells treated with 0.222 ΔM **(Fig.7a)**; for Delta630, 0.074 ΔM treatment could result in significantly less S protein expression compared to the untreated infected cells **(Fig.7b)**; for Omicron, even 0.025 ΔM treatment could result in significantly less S protein expression compared to the untreated infected cells (Fig.7c). Thus, these findings show that HT is exceptionally valuable for anti-SARS-CoV-2 therapy. In summary, this study confirms that HT can inhibit SARS-CoV-2 membrane fusion by double blocking the S protein and TMPRSS2 pathway, thereby making it an effective antagonist for SARS-CoV-2 infection.

**Fig.7.**
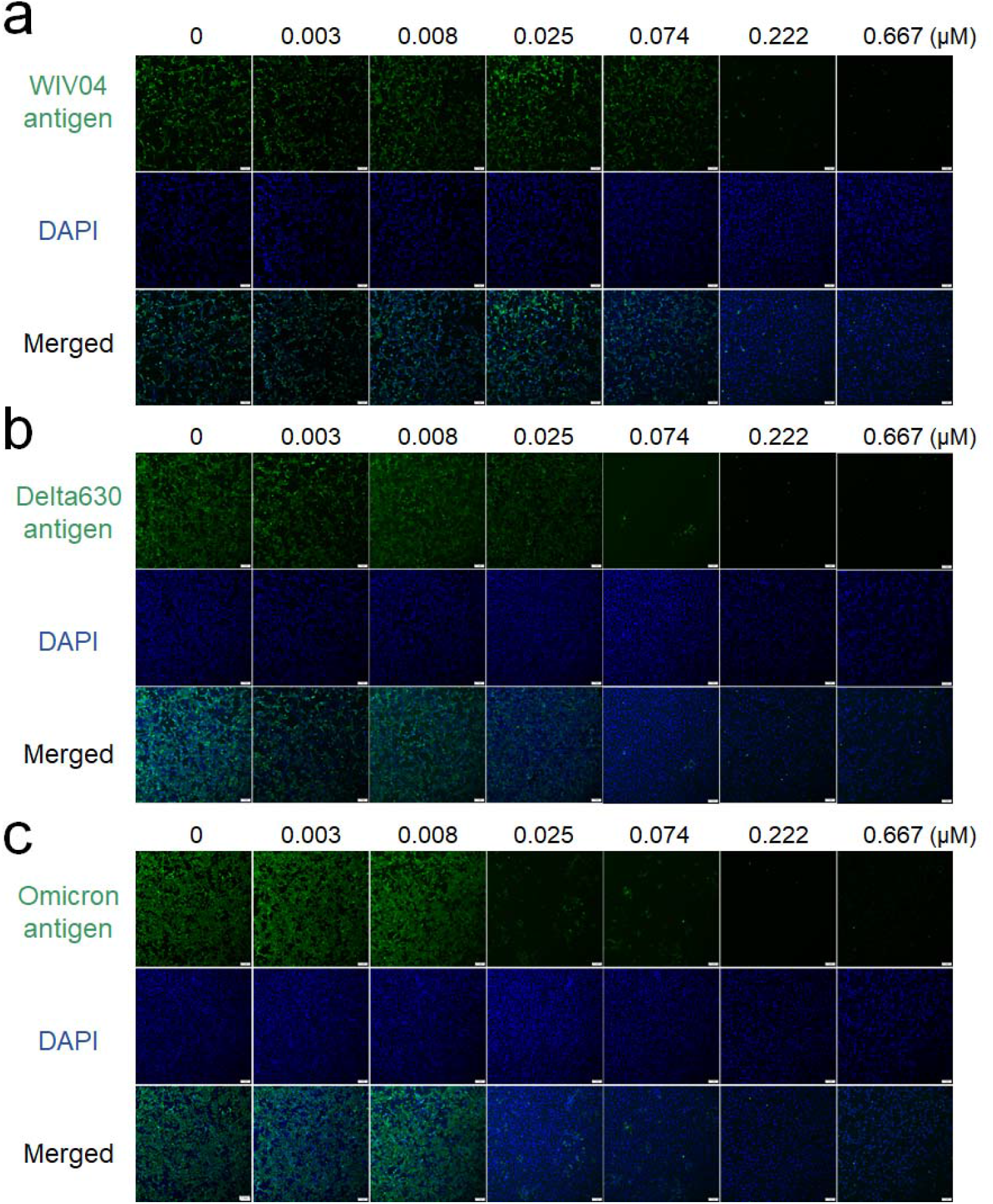
Representative immunofluorescence images of HT significantly inhibited SARS-CoV-2 infection. **a**, Immunofluorescence showed that HT significantly inhibited SARS-CoV-2-WIV04 virus infection in Vero-E6 cells. **b**, Immunofluorescence showed that HT significantly inhibited SARS-CoV-2-Delta 630 virus infection in Vero-E6 cells. **c**, Immunofluorescence showed that HT significantly inhibited SARS-CoV-2-Omicron virus infection in Vero-E6 cells, with a lower inhibiting dose. The scale of the figure is 1 mm.

## Conclusion

Collectively, HT has two potential advantages in the clinical treatment of COVID-19. First, HT mainly targets the S protein and inhibits its ability to bind to ACE2 protein on host cells. Compared with anti-SARS-CoV-2 replication drugs, HT greatly reduces the damage of S protein’s autotoxicity to the organ system. Second, with continuing variation of SARS-CoV-2, the virus will “escape” the recognition of neutralizing antibodies and antibodies produced by inactivated vaccines. However, HT, as a small molecule antagonist, is minimally affected by SARS-CoV-2 variation. At present, the increased infectivity of Omicron reflects that it is stronger membrane fusion effect^24^. TH, as a small molecule antagonist for membrane fusion, is more targeted and can effectively block Omicron infection.

## Acknowledgement

This work was cofounded by National Natural Science Foundation of China (Grant number: 82150201 and 81930096) and the Major Research Development Program of Shaanxi Province (Grant number: 2022ZDXM-SF-03).

